# Non-Invasive Detection of Viral Antibodies Using Oral Flocked Swabs

**DOI:** 10.1101/536227

**Authors:** David J. Speicher, Kathy Luinstra, Emma J. Smith, Santina Castriciano, Marek Smieja

**Affiliations:** Department of Pathology & Molecular Medicine, McMaster University, Ontario, Canada; Department of Laboratory Medicine, St. Joseph’s Healthcare Hamilton, Ontario, Canada; Menzies Health Institute Queensland, Griffith University, Queensland, Australia; M.G. DeGroote Institute for Infectious Disease Research, Department of Biochemistry and Biomedical Sciences, DeGroote School of Medicine, McMaster University, Hamilton, Ontario, Canada; Department of Mathematics and Statistics, University of Guelph, Ontario, Canada; Copan Italia, Brescia, Italy

**Keywords:** oral flocked swabs, saliva, diagnostics, Cytomegalovirus, Epstein-Barr virus, Varicella-Zoster virus, Measles, Mumps

## Abstract

Salivary antibodies are useful in surveillance and vaccination studies. However, low antibody levels and degradation by endonucleases are problematic. Oral flocked swabs are a potential non-invasive alternative to blood for detecting viral antibodies. Serum and saliva collected from 50 healthy volunteers were stored at −80°C; dried swabs at room temperature. Seroprevalence for Cytomegalovirus (CMV), Varicella-Zoster virus (VZV), Epstein-Barr virus (EBV), Measles and Mumps IgG antibodies were determined using commercial ELISAs and processed on an automated platform. For each antibody, swabs correlated well with saliva. For CMV IgG, VZV IgG, and EBV EBNA-1 IgG and VCA IgG, the swab sensitivities compared to serum were 95.8%, 96%, 92.1% and 95.5% respectively. For Measles IgG, swab sensitivity was 84.5%. Mumps IgG displayed poor sensitivity for oral swabs (60.5%) and saliva (68.2%). Specificities for IgG antibodies were 100% for CMV, EBV and Mumps. Specificities for VZV and Measles could not be determined due to seropositive volunteers. As oral flocked swabs correlate well with serum, are easy to self-collect and stable at room temperature further research is warranted.

**Highlights:**

- Oral flocked swabs are an easy, self-collection method for measuring viral antibodies.
- Viral IgG is stable on dried oral flocked swabs for at least two years.
- Oral swabs are highly sensitive for CMV, VZV, and EBV IgG.
- Oral swabs are potentially useful for surveillance and clinical microbiology.

## 1. Introduction

Immunological screening for viral antibodies (primarily IgG) in serum to assess past infection or vaccine immunity is routinely performed via commercial enzyme immunoassays (EIA) on closed platforms. Serum is the gold standard for determining immune status but is invasive to collect. Saliva has considerable diagnostic potential: it is non-invasive, abundant, easily collected, and representative of oral and systemic health. Salivary diagnostics is rapidly emerging, especially defining biomarkers for point-of-care testing of infectious diseases (1). Salivary antibodies are primarily secretory IgA from the salivary glands, while IgG and IgM are derived from serum plasma cells and passively diffused into the oral cavity via gingival crevicular fluid (2, 3). Salivary IgG (sIgG) is systemically representative and strongly correlates with serum levels, but loads are approximately 1:800 that of serum (4, 5). This is problematic for typical closed testing systems that incorporate a 1:100 dilution step. Despite low levels, salivary antibodies are utilized in the U.S. Food and Drug Administration approved OraQuick ADVANCE^®^ Rapid HIV-1/2 Antibody Test (OraSure Technologies, Inc., USA) and OraQuick^®^ HCV Test (OraSure Technologies, Inc.) (1). However, many commercial assays are cost-prohibitive for resource-limited settings.

Saliva collection can be difficult in children and hyposalivators, such as immunocompromised patients. Salivary endonucleases are detrimental, remaining active at −80°C, necessitating special handling, or storage in proteolytic stabilizers unsuitable for antibody preservation (6). To overcome these limitations, procedures for viral antibody detection were optimized on an open commercial platform for dried oral flocked swabs, after room temperature storage. The efficiency of oral flocked swabs to detect viral antibodies has yet to be determined. Our method was initially optimized for Cytomegalovirus (CMV) IgG due to its importance in hematopoietic stem cell, solid organ, and haploidentical transplantations as well as prenatal patients (7, 8). We then assessed the procedures’ potential application to detect Varicella-Zoster virus (VZV), Epstein-Barr virus (EBV), Measles and Mumps IgG.

## 2. Materials and Methods

### 2.1. Study Population

Following Hamilton Integrated Research Ethics Board (HiREB #14-658) approval and written informed consent, two oral swabs, unstimulated saliva, and blood were collected. Optimisation of pre-analytic and analytic procedures was performed on 10 healthy volunteers with known CMV seropositivity (5 positive, 5 negative), and expanded to 50 healthy volunteers for the diagnostic accuracy study. Laboratory staff (15 males:35 females) from St. Joseph’s Healthcare Hamilton with an average age of 43.4 years (range: 18-65 years) voluntarily provided all sample types, except for one who could not produce a saliva sample. Two oral swabs (FLOQSwabs^®^ #520C, Copan Italia S.p.A., Brescia, Italy) were collected consecutively by moistening the flocked swab on the tongue and then rotating between the gums and cheek three to five times. Swabs were then dried for an hour inside a biosafety cabinet and stored inverted in a microcentrifuge tube at room temperature prior to elution and at −20°C following elution. Whilst circadian rhythm was not accounted for as swabs were collected at times convenient for the volunteer, participants were asked to refrain from eating or drinking 60-minutes prior to collection. Cell-free unstimulated saliva was collected by expectorating 2-5mL into a sterile 50mL Falcon tube, centrifuging at 2,800 *x g* for 10-minutes and aspirating the supernatant (9). Supernatant was aliquoted into 1mL portions and stored at −80C; the cell pellet was discarded. Serum was obtained from a 5mL blood collection via venipuncture, allowed to clot, centrifuged at 3,000 *x g* for 10-minutes, and stored at −80°C.

### 2.2. Development and Optimisation of pre-analytic and analytic procedures

The first optimisation to determine the optimal dilution for testing oral swabs was performed with a CMV IgG EIA on the ThunderBolt^®^ ELISA Analyzer; both from Gold Standard Diagnostics (GSDx, Davis, CA, USA). To elute viral antibodies, 250μL PBS was added to a dried swab head, vortexed for 30-seconds, incubated at room temperature for 10-minutes, and centrifuged at 14,000 *x g* for one minute, before discarding the swab. Serial dilutions (2-fold serially from neat to 1:16) were prepared in PBS and 100μL tested on the ThunderBolt^®^ ELISA Analyzer as per manufacturer’s instructions. Repeated measures one-way ANOVA was performed in R3.5.0 in combination with polynomial contrasts to assess the nature and significance of the relationship between dilutions and optical density (O.D.) values (10).

The second optimisation determined the effect of tube shape and elution volume on O.D. values and status. Various volumes of PBS (150μL, 200μL, and 250μL) were added to swabs stored in two shapes of microcentrifuge tubes: 2.0mL flat-bottomed, screw cap tubes (SCT-200-Y, Axygen Scientific, Union City, CA, USA) and 1.5mL conical microcentrifuge tubes (MCT-150-C, Axygen Scientific). The volume of PBS recovered was measured for each. To assess the relationship between O.D. values and both tube shape and volume, a linear mixed effects model was fit. O.D. values were treated as the response and volume added and tube type were considered as the main effects. As sample manipulations could affect positivity only positive patients were considered.

The third optimization determined the effect pelleted buccal cells had on O.D. values and positivity of five salivopositive and five salivonegative samples using three methods: 1. Pellet, measuring the supernatant O.D. values with undisturbed pelleted buccal cells at the bottom of the tube; 2. Supernatant, measuring the supernatant O.D. values after transfer to a new tube without disturbing the pellet; 3. Resuspended; measuring the O.D. values after complete resuspension of the buccal cells. To determine the relationship between resuspension methods, an initial ANOVA followed by paired testing was used. Further pairwise comparisons between O.D. values of the three methods (pellet, supernatant, and resuspended) were constructed using paired t-tests.

The fourth optimisation determined the stability of viral antibodies in dried flocked swabs over time by measuring O.D. values from five salivopositive samples at baseline and after two months stored at room temperature. Paired t-tests were used to compare average O.D. values at both time points.

### 2.3. Diagnostic Accuracy Study

A cross-sectional diagnostic accuracy study was performed to compare oral flocked swabs and unstimulated saliva versus serum as a reference standard, for the detection of various viral-specific IgG antibodies. As results were similar between two consecutively collected swabs (data not shown) swab eluates were pooled to facilitate automated testing of multiple analytes. The assays utilized included: CMV IgG EIA, EBV EBNA IgG EIA, EBV VCA IgG EIA, Measles IgG EIA, and Mumps IgG EIA from GSDx; RIDASCREEN^®^ VZV IgG (K5621), RIDASCREEN^®^ Measles IgG (K5421), and RIDASCREEN^®^ Mumps IgG (K5521) from R-Biopharm AG (Darmstadt, Germany). These commercial EIAs are optimised for serum but were used off-label for oral fluids. All assays were performed as per manufacturer’s protocol using 100μL 1:100 sera, but 100μL undiluted oral fluid. For each assay, the mean O.D. and standard deviation (SD) were calculated from swabs and saliva specimens corresponding to “true-negative” subjects, i.e. serum test negative subjects. We then defined cut-off values as: Non-reactive if less than two SDs above the average O.D. values; Reactive if greater than 3 SDs above the average O.D.; all other O.D. values considered indeterminate. As all samples were seropositive for VZV and measles the cut-off O.D. values for these analytes were extrapolated from the cut-off O.D. values of other assays from the same manufacturer. For each, the diagnostic test accuracy (sensitivity, specificity, positive predictive value (PPV), negative predictive value (NPV), and overall accuracy), and misclassification rates were determined. Kappa statistics were calculated in a pairwise fashion to quantify the agreement beyond chance between oral swabs, unstimulated saliva, and serum for all antibodies of interest.

## 3. Results

### 3.1. Optimisation of pre-analytic procedures

To determine the effect of sample dilutions, O.D. values were measured at five, two-fold dilution points. Dilutions significantly decrease O.D. values (*p*=0.013) and affected positivity: 4-fold and 8-fold dilutions yielded 2/5 (40%) and 4/5 (80%) false non-reactive samples, respectively (Figure 1). Pairwise comparison showed that dilutions significantly decreased O.D. values in a linear relationship (F=17.758, *p*=0.014). Therefore, swabs were used undiluted to ensure that weakly reactive samples did not become falsely non-reactive.

**Figure 1.**
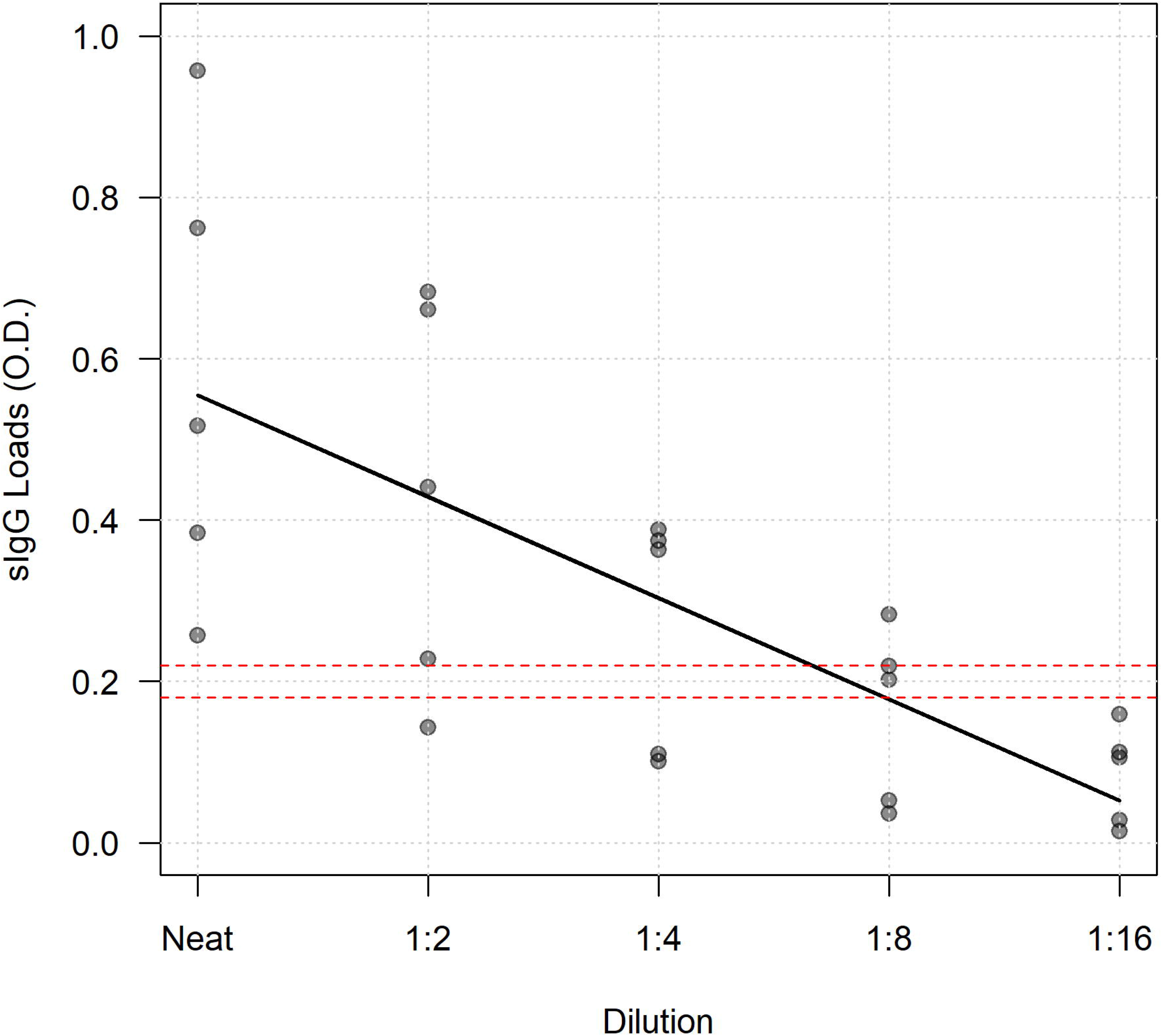
Two-fold dilutions of the swab eluate in PBS with detection for CMV IgG. Dashed lines indicate the reactive (upper; O.D. = 0.291) and non-reactive (lower; O.D. = 0.222) cut-off O.D. values.

To determine the effect of tube shape and establish the optimal elution volume, O.D. values were measured at three volumes of PBS and two tube shape combinations. The conical tube shape caused eluate to be reabsorbed by the swab, significantly reduced recovery volume (conical vs flat-bottomed: 60.66 ± 4.38% vs 98.63 ± 0.78%; *p*=0.017), O.D. values (*p*=0.003), and potentially affected the positivity of weakly reactive samples (Figure 2). The O.D. values slightly decreased with increased volume added (150μL to 250μL), but this difference was not significant (*p*=0.184) and did not affect positivity. In the non-reactive samples, the average O.D. values of the flat-bottomed tubes (0.080 ± 0.008) were slightly higher than the conical tubes (0.059 ± 0.021), but this difference was not significant (*p*=0.323). Therefore, subsequent testing was performed by eluting with 250μL PBS in flat-bottomed tubes to facilitate two tests per swab.

**Figure 2.**
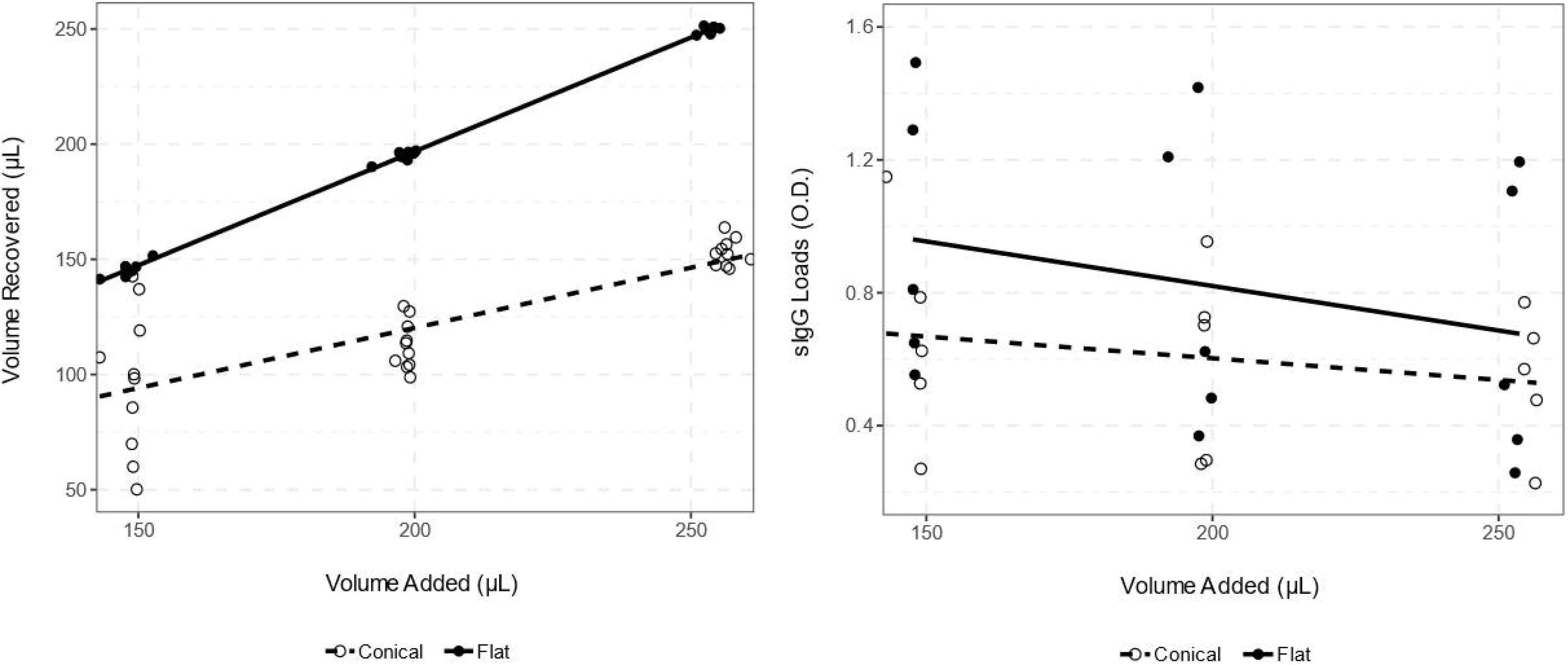
Graphical representation showing how flat-bottom (solid circles and line) and conical (empty circles and dashed line) tube shape affects (A) volume recovered and (B) CMV sIgG loads.

To determine the most efficient sample handling procedure to yield the highest O.D. values, three methods were evaluated. In the reactive samples, there was little difference between the O.D. values: undisturbed pellet (0.688 ± 0.434); supernatant (0.545 ± 0.222); resuspended pellet (0.531 ± 0.214). One sample was a weak reactive in the pellet method (O.D. 0.258) but became indeterminate in the supernatant (O.D. 0.214) and resuspended pellet (O.D. 0.197) method. Pairwise comparison revealed no significant differences in O.D. values between the sample handing methods (pellet: supernatant, *p*=0.312; pellet:resuspended, *p*=0.288; supernatant:resuspended, *p*=0.515), but fractioning the sample could produce an indeterminate result from a weakly reactive sample and was more labour intensive. Therefore, subsequent testing was performed using the whole sample following centrifugation.

To determine the stability of oral swab collections at room temperature, O.D. values were measured from dried oral swabs the day of collection and after two months of storage at room temperature. The average O.D. value did not significantly differ between baseline (O.D. 0.575 ± 0.284) and two months (O.D. 0.568 ± 0.188). The mean difference in O.D. values was 0.008 (p=0.946). No change is salivopositivity was observed.

### 3.2. Diagnostic Accuracy Study

To determine the correlation between oral flocked swabs, unstimulated saliva and serum a cross-sectional diagnostic accuracy study was conducted on 50 volunteers for CMV, VZV, EBV EBNA-1 and VCA, Measles, and Mumps IgG. For CMV IgG, the seropositivity using serum was 24/50 (48.0%). The cut-off O.D. values for swabs and saliva were non-reactive <0.179 and reactive >0.221; and non-reactive <0.220 and reactive >0.253, respectively (Table 1). Using the new cut-off O.D. values the sensitivity of swabs and saliva were 23/24 (95.8%; 95% CI: 78.1%, 100%) and 24/24 (100%; 95% CI: 83.7%, 100%), respectively (Table 2). Specificity of both swabs and saliva were 100%. One swab was indeterminate. The agreement beyond chance was very good between oral swabs and both serum (K=0.88; 95% CI: 0.76, 1.000) and saliva (K=0.85; 95% CI: 0.71, 1.00), and perfect between saliva and serum (K=1.00; 95% CI: 0.86, 1.00) (Figure 3).

**Figure 3.**
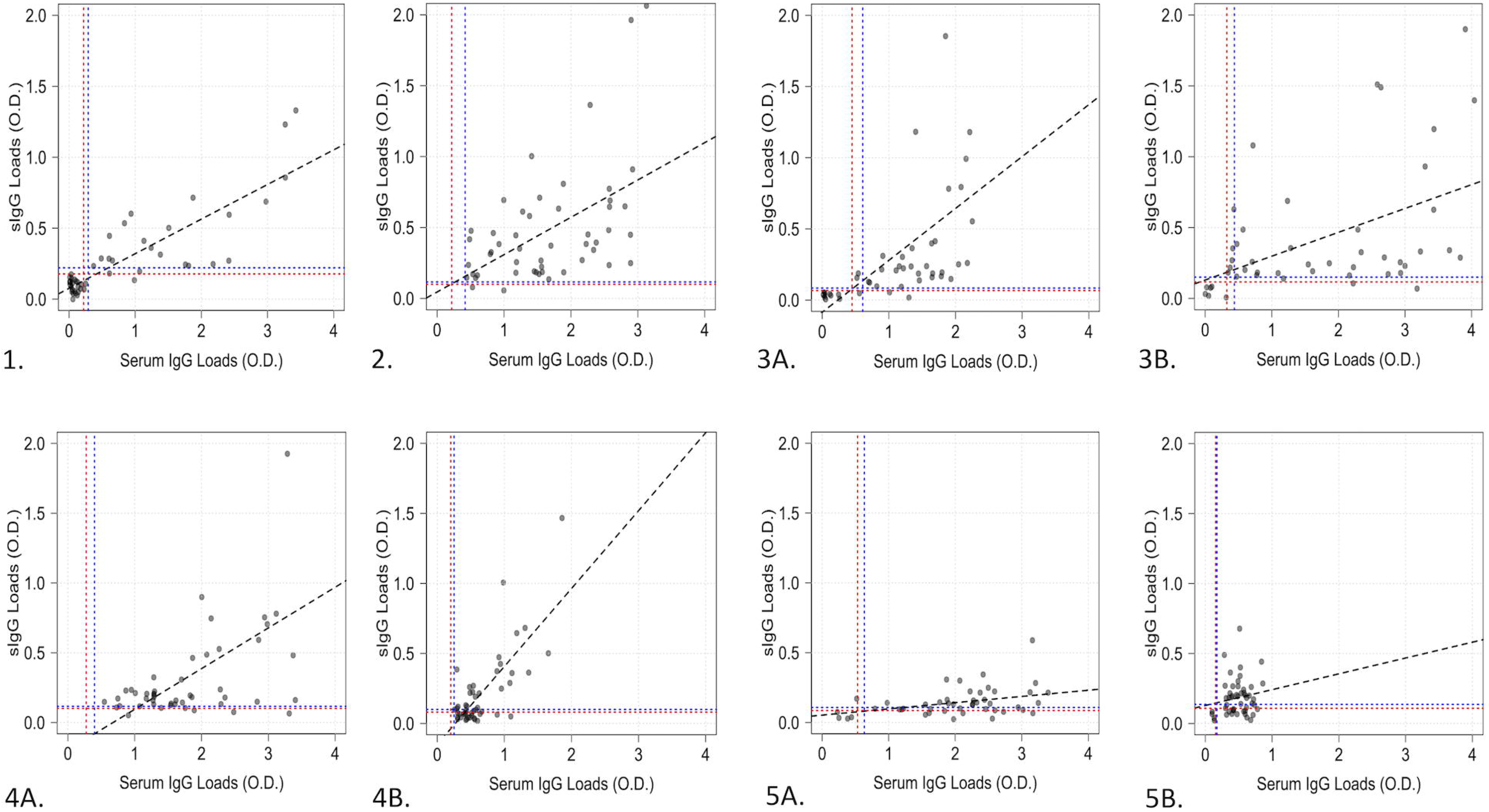
Comparison of viral-specific sIgG in paired flocked swab versus serum samples from 50 healthy volunteers for the following targets: 1. CMV; 2. VZV; 3. EBV; 4. Measles; 5. Mumps. For EBV both (a) EBNA-1 and (b) VCA were tested. For Measles and Mumps assays from (a) Gold Standard Diagnostics and (b) R-Biopharm AG were investigated. Positive cut-off values (dotted blue line) and negative cut-off values (dotted red line) were determined for each assay.

**Table 1.**
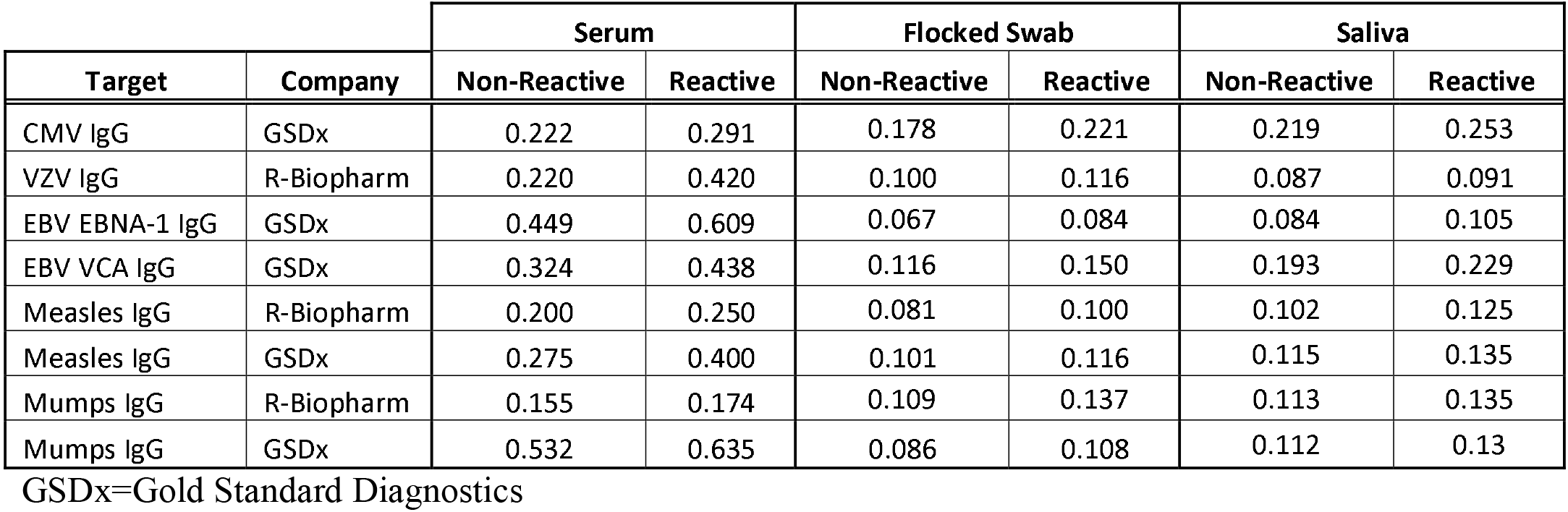
Cut-off O.D. values for viral-specific IgG assays. For each assay the cut-off O.D. values were determined as follows from the average O.D. values of all negative samples: Non-reactive = O.D. < mean (non-reactive) + 2 St. Dev; Indeterminate = > mean (non-reactive) + 2 St Dev but < mean (non-reactive) + 3 St Dev; Reactive = O.D. > mean (non-reactive) + 3 St Dev.

**Table 2.** Diagnostic accuracy for viral-specific IgG assays in swabs and saliva compared to sera. For all samples were seropositive for VZV and Measles so the specificity could not be determined.

|  | Cytomegalovirus |  | Varicella Zoster Virus |  | Epstein-Barr Virus |  |  |  | Measles |  |  |  | Mumps |  |  |  |
| --- | --- | --- | --- | --- | --- | --- | --- | --- | --- | --- | --- | --- | --- | --- | --- | --- |
| Assay | GSDx CMV IgG |  | R-Bio VZV IgG |  | GSDx EBNA-1 IgG |  | GSDx VCA IgG |  | GSDx MeV IgG |  | R-Bio MeV IgG |  | GSDx MuV IgG |  | R-Bio MuV IgG |  |
| Specimen | Swabs | Saliva | Swabs | Saliva | Swabs | Saliva | Swabs | Saliva | Swabs | Saliva | Swabs | Saliva | Swabs | Saliva | Swabs | Saliva |
| Sample (n=) | 50 | 49 | 50 | 49 | 50 | 49 | 50 | 49 | 50 | 49 | 50 | 49 | 50 | 49 | 50 | 49 |
| Positives* | 23/24 | 24/24 | 48/50 | 46/49 | 35/38 | 34/38 | 42/44 | 42/44 | 41/50 | 46/50 | 24/50 | 34/50 | 27/45 | 31/45 | 28/46 | 33/46 |
| Sensitivity | 95.8% | 100.0% | 96.0% | 93.9% | 92.1% | 91.9% | 95.5% | 97.7% | 85.4% | 93.9% | 49.0% | 69.4% | 60.5% | 68.2% | 62.2% | 73.3% |
| Specificity | 100% | 100% | ND | ND | 100% | 100% | 100% | 100% | ND | ND | ND | ND | 80.0% | 80.0% | 100.0% | 100.0% |
| PPV | 100% | 100% | 100% | 100% | 100% | 100% | 100% | 100% | 100% | 100% | 100% | 100% | 96.3% | 96.8% | 100.0% | 100.0% |
| NPV | 96.3% | 100% | ND | ND | 80.0% | 80.0% | 75.0% | 85.7% | ND | ND | ND | ND | 19.0% | 22.2% | 19.0% | 25.0% |
| Accuracy | 98.0% | 100% | 96.0% | 93.9% | 94.0% | 93.9% | 96.0% | 98.0% | 85.4% | 93.9% | 49.0% | 69.4% | 62.5% | 69.4% | 65.3% | 75.5% |
| Misclassification | 4.0% | 0.0% | 4.0% | 6.1% | 6.0% | 6.1% | 4.0% | 2.0% | 10.0% | 6.1% | 50.0% | 30.6% | 40.9% | 33.3% | 37.8% | 26.7% |
| Agreement (K) | 0.88 | 0.85 | 0.00 | 0.00 | 0.85 | 0.85 | 0.83 | 0.91 | 0.00 | 0.00 | 0.00 | 0.00 | 0.20 | 0.25 | 0.21 | 0.32 |
| Linearity (Adj. R<sup>2</sup>) | 0.713 | 0.622 | 0.248 | 0.218 | 0.315 | 0.178 | 0.232 | 0.266 | 0.308 | 0.351 | 0.554 | 0.316 | 0.142 | 0.221 | 0.005 | 0.017 |

\*# of swab or saliva positives/# serum positive samples; PPV = Positive Predictive Value; NPV = Negative Predictive Value; GSDx = Gold Standard
Diagnostics; R-Bio = R-Biopharm AG; ND = Not Determined.

For VZV IgG, all participants were seropositive. There was excellent correlation between both swabs and saliva to serum: 48/50 (96.0%; 95% CI: 85.7%, 99.7%) and 46/49 (93.9%; 95% CI: 82.9%, 98.5%), respectively. As there were no seronegative participants specificity could not be determined and Cohen’s kappa coefficient was poor for both sera to swab and saliva (K=0), and fair for swabs vs saliva (K=0.37; 95% CI: −0.189, 0.928).

For EBV, fewer participants were seropositive for EBNA-1 IgG than VCA IgG: 38/50 (76.0%) and 44/50 (88.0%), respectively. The sensitivity of swabs and saliva were comparable for both EBNA-1 IgG [swabs: 35/38 (92.1%; 95% CI: 78.5%, 98.0%); saliva: 34/38 (91.9%; 95% CI: 75.3%, 96.4%)] and VCA IgG [swabs: 42/44 (95.5%; 95% CI: 84.0%, 99.6%); saliva: 42/43 (97.7%; 95% CI: 86.8%, 100%)]. However, sensitivity for the composite measure of EBNA-1 IgG *or* VCA IgG equated to 43/44 (97.7%; 95% CI: 87.1%, 100%) and 44/44 (100%; 95% CI: 90.4%, 100%) for swabs and saliva, respectively. The agreement for EBNA-1 IgG was very good for both swab (K = 0.85; 95% CI: 0.68, 1.00) and saliva (K=0.85; 95% CI: 0.68, 1.00) compared to serum, and perfect between swabs and saliva (K=1. 000; 95% CI: 0.90, 1.00). The agreement for VCA IgG was very good for both swab (K=0.83; 95% CI: 0.61, 1.000) and saliva (K=0.91; 95% CI: 0.74, 1.00) compared to serum, and good between swabs and saliva (K=0.76; 95% CI: 0.50, 1.00).

For Measles and Mumps IgG the correlation between oral swabs, unstimulated saliva and serum as well as between two EIAs was determined. For Measles IgG, all participants were seropositive by both assays. The sensitivity of swabs and saliva was much higher for the GSDx assays for both swabs [41/48 (85.4%; 95% CI: 72.5%, 93.1%) vs 24/49 (48.9%; 95% CI: 35.6%, 62.5%)] and saliva [46/49 (93.9%; 95% CI: 82.9%, 98.5%) vs 34/49 (69.4%; 95% CI: 55.4%, 80.6%)]. However, the agreement for both assays between swab and saliva compared to sera was poor (K=0), and poor for swab compared to saliva (GSDx: K=0.24; 95% CI: −0.08, 0.56; R-Biopharm: K=0.27; 95% CI: −0.09, 0.46). Further analysis showed the R-Biopharm assay had a much larger misclassification rate for both swab [25/50 (50.0%; 95% CI: 36.6%, 63.4%) vs 5/50 (10.0%; 95% CI: 3.9%, 21.8%)] and saliva [15/49 (30.6%; 95% CI: 19.5%, 44.6%) vs 3/49 (6.1%; 95% CI: 2.1%, 16.5%)], suggesting that the GSDx assay performs better for measuring measles salivopositivity.

For Mumps IgG, both assays produced comparable seroprevalence of 45/50 (90.0%) and 46/50 (92.0%) for the GSDx and R-Biopharm assays, respectively. The sensitivity of both assays was poor for both swabs [GSDx: 26/43 (60.5%; 95% CI: 45.6%, 73.7%); R-Biopharm: 28/45 (62.2%; 95% CI: 47.6%, 74.9%)] and saliva [GSDx: 30/44 (68.2%; 95% CI: 53.4%, 80.1%); R-Biopharm: 33/45 (73.3%; 95% CI: 58.4%, 84.2%)].…(11). The agreement was only fair for both assays between swab (GSDx: K=0.20; 95% CI: 0.00, 0.40; R-Biopharm: K=0.21; 95% CI: 0.03-0.40) and saliva (GSDx: K=0.25; 95% CI: 0.02, 0.47; R-Biopharm: K=0.32; 95% CI: 0.07, 0.56) compared to sera, as well as swabs compared to saliva (GSDx: K=0.25; 95% CI: 0.03, 0.47; R-Biopharm: K=0.38; 95% CI: 0.16, 0.60). Further analysis showed that both assays had a large misclassification rate for both swab [GSDx: 18/44 (40.9%; 95% CI: 27.7%, 55.6%); R-Biopharm: 17/45 (37.8%; 95% CI: 25.1%, 52.4%)] and saliva [GSDx: 15/45 (33.3%; 95% CI: 21.3%, 48.0%); R-Biopharm: 12/45 (26.7%; 95% CI: 15.8%, 41.2%)], suggesting that neither assay is ideal for measuring mumps salivopositivity.

## 4. Discussion

Oral fluids contain IgG profiles highly similar to those in serum for a range of antigens and diseases regardless of anatomical location (12). Their non-invasive collection is utilized in several point-of-care assays to detect viral infections and immunity. To avoid degradation by salivary endonucleases, diagnostic assays either utilize unadulterated gingival crevicular fluid or whole saliva either chilled immediately following collection and stored frozen or stored in biological stabilizers (4, 6). Studies for viral IgG utilize the Oracol saliva collection system (Malvern Medical Developments, UK), which collects 1mL saliva into 1mL transport medium allowing transportation at room temperature (13, 14). However, as samples must be stored frozen for long-term preservation this is not ideal for field studies and resource-limited settings (15). Therefore, as FLOQSwabs^®^ are routinely used for bacteriology and molecular microbiology (16, 17), we developed a simple pre-analytic and analytic method for detecting viral sIgG on an open commercial platform using dried FLOQSwabs^®^ that can be stored at room temperature for up to 2-months. This was not an epidemiological study, but our method is applicable to point-of-care or home testing, surveillance, clinical epidemiology and clinical microbiology.

Our protocol was developed on a convenient cohort of healthy volunteers and appears promising as swabs can be self-collected and diagnostic accuracy is comparable to saliva and serum for most viruses tested. The similarity between swabs and saliva is due to cell-free viral IgG secreted via the gingival crevicular fluid into the mouth and absorbed onto the swab regardless of oral location swabbed. As swab absorption volume is much lower than the volume of saliva in the mouth it is possible to collect multiple swabs consecutively at any time of day without a reduction in positivity. Storing swabs dried maintains sample integrity and restricts activity of salivary endonucleases for at least 2-months post-collection. Ideally, swabs could be collected and shipped, but further investigation is needed to assess drying time on sample stability. Nevertheless, a few procedural steps must be heeded. Whilst cut-off O.D. values vary between assays, platforms and/or cohort tested, it is essential to elute swabs into flat-bottomed tubes to prevent reabsorption and then test undiluted eluate. Elution with 250μL PBS permits each sample to be tested twice. Further work is needed to (i) optimize elution and testing volumes to maximize testing and increase O.D. values to resolve indeterminate samples, (ii) determine sample integrity by comparing the absolute amount of human IgG, and (iii) compare manual testing vs automation to determine the robustness of the protocol.

Our swab protocol was initially developed for CMV due to interest from blood banks and transplant programs. CMV is readily shed in saliva, the viral load is 100-fold higher, and the limit of detection is 10-fold lower in saliva collected with a sterile swab than urine (18–20). CMV antibody profiles in serum also strongly correlate with CMV infection and oral shedding (8, 21, 22), and, based on our study, CMV IgG can be detected accurately in oral swabs. Swabs displayed similar diagnostic accuracy to saliva and serum. The overall O.D. values in oral fluids was lower than serum with swabs slightly lower than saliva but displayed a linear comparison in O.D. values between serum and oral fluids. The slightly lower swab O.D. values compared to saliva results from a fraction of whole saliva diluted in eluate. The one swab that was equivocal was strongly positive in sera and saliva suggesting further optimisation is warranted. Whilst our protocol was developed for CMV IgG in healthy volunteers and may not be reflective of assay performance in hospital or immunocompromised patient populations, it worked well for herpesviruses due to maintained immunity from latent infections, but less than optimal for measles and mumps possibly due to waning vaccine-induced immunity.

Few publications examined salivary IgG for VZV and EBV. In the only study on VZV IgG in oral fluids, Talukder *et al*. examined 1,092 participants and reported a sensitivity and specificity of 93% and 95.7%, respectively (14). The sensitivity is comparable to saliva (93.9%) and swabs (96.0%) in our study, but our sample size was insufficient to determine assay specificity. In a hospital laboratory volunteer study, virtually all employees have either received screening and vaccination for vaccine-preventable illnesses or are old enough to have natural immunity. Based on the diagnostic accuracy our procedure could be studied as a screening method for preschool children susceptible to chicken pox.

Epidemiological screening for EBV, the aetiological agent of infectious mononucleosis and nasopharyngeal carcinoma, is performed by screening for EBV VCA IgM, VCA IgG, and EBNA-1 IgG via IFA or EIAs to distinguish acute from past infection (23). Both Vyse *et al*. and Crowcroft *et al*. utilized oral fluids to determine EBV immune status (24, 25). Vyse *et al*. used a ‘G’ antibody capture radioimmunoassay for EBV VCA IgG and reported a sensitivity and specificity of 93.5% and 100%, respectively (25). Their assay was less sensitive than IFA as samples with a total IgG <2 mg/L yielded false positive reactions due to the monoclonal antibody binding non-specifically to unsaturated anti-human IgG on the solid phase. Although we didn’t examine the total IgG in our samples, our protocol produced higher sensitivities for both swabs and saliva with no known false positives. As 5% of people do not produce EBNA-1 IgG after EBV infection we validated both EBV EBNA-1 IgG and VCA IgG (26). Combining both assays our protocol yielded a sensitivity of 97.7% and 100% for swabs and saliva, respectively, and may be useful for surveillance studies and could be preferred over serum heterophile antibody testing for acute infectious mononucleosis work-up.

As Measles and Mumps are vaccine-preventable infections several studies have used oral fluids as a non-invasive alternative for monitoring the efficacy of vaccination programs. For Measles studies have used off-label either the commercialized Measles IgG Capture EIA (Microimmune Ltd., UK), and the Enzygnost^®^ Anti-Measles Virus/IgG (Siemens Health Care Diagnostics GmbH, Germany) with mixed results (5). Whilst Hayford *et al*., reported that oral fluids are not suitable to detect immunity for Measles due to poor sensitivity (60.2%) and specificity (75.7%) (27, 28) others reported sensitivity of 90.0-92.0% and a specificity of 77.8-100% using the Microimmune Ltd. assay (29–32). Our study also produced mixed results between the GSDx and R-Biopharm assays. Whilst the R-Biopharm assay produced poorly, the GSDx assay was comparable to the Microimmune Ltd. assay. Further optimization is required to increase the sensitivity of our assay, but in its present stage may be adequate for epidemiological studies. If the efficacy of vaccination is essential, confirmatory testing on serum should be performed for non-reactive oral samples.

For Mumps, Vainio *et al*., assessed the Mumps IgG Capture EIA (Microimmune Ltd.) and reported low detection of Mumps IgG in oral samples (76% sensitivity) and recommended that the Microimmune assay not be used for surveillance studies (33). In our study, both the GSDx and R-Biopharm assays produced similarly low sensitivities for both swabs and saliva, but possibly for different reasons. The R-Biopharm assay produced low O.D. values for both oral fluids and serum suggestive of an issue with the assay. The GSDx assay yielded a range of O.D. values for serum similar in range to the other assays, but much lower (<0.5) for swabs. Whilst it is possible that Mumps IgG is not excreted into the mouth, vaccine-derived immunity to Mumps also wanes especially into adulthood and maybe too low to detect in oral fluids (34).

Whilst the use of oral fluids is not novel, we developed a simple non-invasive protocol for detecting viral IgG using oral flocked swabs. Based on the high sensitivity and excellent correlation with serum, oral samples are ideal for CMV, EBV, and VZV, adequate for Measles, but poor for Mumps. As samples can be stored dried for a few months, they are a viable option for home and field collection. Future work should investigate the utility of oral swabs for viral IgM as well as Rubella. Studies using oral fluids for Rubella IgM look promising with 79-96.9% sensitivity and 90-100% specificity (32, 35). Nonetheless, oral swabs appear promising for surveillance studies, transplant screening programs, and clinical microbiology.

## 5. CRediT Author Statement

David J. Speicher: Conceptualization, Investigation, Writing - Original Draft; Kathy Luinstra: Methodology, Writing - Review & Editing; Emma J. Smith: Formal analysis; Santina Castriciano: Conceptualization, Resources, Funding acquisition; Marek Smieja: Supervision

## 6. Acknowledgements

This study was supported in part by an award from The Research Institute of St. Joe’s Hamilton. We are grateful for the donation of FLOQSwabs^®^ (#520C) from Copan Italia, and commercial EIA kits from Gold Standard Diagnostics and R-Biopharm AG. We thank Gold Standard Diagnostics for lending the ThunderBolt^®^ ELISA Analyzer. This study was based on the initial CMV IgG testing by Dr Milena Furione and presented at the Clinical Virology Symposium (CVS 2016; Poster #230). Santina Castriciano is an employee of Copan Italia. The other authors have no financial or other conflicts of interest to declare. Whilst this manuscript was shared with Copan Italia and Gold Standard Diagnostics prior to submission these collaborators had no influence on the data analysis or publication. We are thankful for the support from Jalees Nasir for the statistical help. We are grateful to the many volunteers at St. Joseph’s Healthcare Hamilton who provided sera, saliva, and oral swabs for testing.

